# Immune organization in sentinel lymph nodes of melanoma patients is prognostic of distant metastases

**DOI:** 10.1101/2024.11.24.625041

**Authors:** Yael Amitay, Idan Milo, Tal Keidar Haran, Saif Deis, Guy Truzman, Ofer Elhanani, Tomer-Meir Salame, Michal Azimov, Ilan Stein, Jonathan E. Cohen, Michal Lotem, Eli Pikarsky, Leeat Keren

## Abstract

Sentinel lymph node (sLN) biopsy is part of melanoma staging, as involved LNs indicate a higher risk of recurrence. However, how the sLN is shaped by the tumor and reciprocally affects metastatic progression is poorly understood. Here, we mapped immune organization in involved and non-involved sLNs of 69 melanoma patients using high-resolution spatial proteomics, spatial transcriptomics and deep learning, leveraging the data for prognostic evaluation. In patients with involved LNs, a robust T cell response correlated with absence of recurrence. In non-involved LNs, protection from metastases was linked to expanded sinuses colocalized with plasmablasts whereas CCR7+ cells and Tregs correlated with future development of distant metastases. We trained a model that predicts development of distant metastases with 79% and 93% AUC for non-metastatic and metastatic LNs, respectively. Our findings reveal conserved immune patterns in sLNs that prospectively identify patients at risk for metastatic disease and may aid in therapeutic decisions.

## Introduction

Malignant melanoma ranks as the fifth most prevalent cancer in the Western world, with its frequency steadily increasing. Approximately 85% of melanomas are identified at a very early stage amenable to surgical resection, resulting in a favorable prognosis with a five-year survival rate exceeding 98%. However, metastatic melanoma is deadly, responsible for over 80% of fatalities associated with all skin cancers^1,2^. This discrepancy in prognosis between localized and disseminated disease presents a significant clinical dilemma. Early stage melanoma affords a window of opportunity for potentially curative therapy, thus one could advocate for treating primary tumors aggressively, as if they were metastatic, utilizing approaches such as immunotherapy. Conversely, such a strategy risks overtreating numerous patients, leading to increased morbidity and mortality and severe chronic side effects^3^.

The current clinical workflow to establish if the cancer has spread is to focus on the sentinel lymph nodes (sLNs). The sLN is the first lymph node (or group of nodes) connected to a primary tumor via lymphatic vessels, and thus the one to which cancer cells are most likely to spread first. During cancer staging, the sLN is identified, removed, and examined to determine whether cancer cells are present^4^. If metastases are found in the sLN, the cancer will be classified as stage III and the patient will likely be treated systemically, usually with targeted therapy or immunotherapy. Conversely, negative sLNs confer a stage II tumor, and, in the absence of high risk features of the primary tumor, this allows a surveillance only approach^5^. Although sLN involvement is a helpful prognostic factor and part of the routine guidelines for melanoma staging, 44% of patients with positive sLNs will not experience disease relapse^6^. Moreover, 10-25% of patients with negative sLNs will proceed to develop recurrent metastases within three years of diagnosis^6,7^. As the indications for systemic treatment following local excision are being expanded to include patients with stage II in addition to stage III disease^8^, tools to better identify the subgroups of patients who stand to benefit from these potentially toxic and expensive therapies are needed.

LNs are not simply waystations for transiting tumor cells but serve as hubs of anti-tumor immunity. The sLN, as the first responding LN to the tumor, is shaped by tumors, and has the potential to reciprocally target the tumor. In the course of the tumor immunity cycle, tumor-derived antigens are drained to the LNs, or actively trafficked to the LNs by dendritic cells (DCs) and presented to T cells to elicit an anti-tumor immune response. Presentation of tumor antigens by DCs in the LNs has been shown to drive anti-tumor immunity^9–11^, and the success of immunotherapy has been shown to require activation of immune responses in LNs in addition to the immediate tumor microenvironment^12–14^. However, recent studies have shown that the presence of metastases in tumor-draining LNs are associated with immunosuppressive cellular niches and impaired anti-tumor responses^15–18^. Moreover, work in mice has suggested that the presence of metastases in tumor-draining LNs, in addition to its ability to promote local immune tolerance, may facilitate further metastatic spread by promoting systemic immune tolerance ^19^. Thus, murine models have shown that LNs may harbor conflicting roles, both suppressing and enabling tumor spreading. Knowledge of LN involvement in the metastatic process in human melanoma is even more limited.

We hypothesized that sLNs may play both a pro-tumorigenic and an anti-tumorigenic role, and that they may assume different roles in different patients and along different stages of the disease. Moreover, we hypothesized that this role may influence systemic immunity and therefore be predictive for the development of distant metastases, thus addressing an urgent clinical need. To this end, we assembled a retrospective cohort of negative and metastatic sLNs from 69 melanoma patients, ensuring that approximately half of the individuals in both groups developed distant metastases. Using high-resolution spatial multiplexed imaging, we analyzed 39 proteins with MIBI-TOF ^20^ and 960 mRNAs with CosMX ^21^ to create an atlas of cell types and states within sLNs. We developed a deep learning method to map unique microenvironments in these LNs, revealing both known LN structures and novel microenvironments including one that was enriched with Tregs, CD8 T cells, and IDO-1^+^ dendritic cells. We found that although tumor cells colonizing LNs are in immune hubs, they foster a local microenvironment that is enriched with neutrophils, macrophages and exhausted CD8 T cells. In patients with metastatic sLNs, protection from distant metastases correlated with increased activation of CD8 T cells in the LN metastasis, and Tregs in the follicles, potentially indicating a robust immune response. In patients with negative sLNs, two distinct patterns emerged: those who did not develop distant metastases exhibited an expanded sinus with colocalizing plasmablasts, while those who developed distant metastases had increased Tregs and an influx of CCR7^+^ immune cells in the T zone, potentially promoting tolerance. By integrating cell type composition, phenotype, and microenvironmental data, we achieved an area under the curve (AUC) of 79% and 93% in predicting recurrent metastases for negative and metastatic LNs, respectively. Our findings underscore the complex interplay between pro- and anti-tumorigenic immune responses in sLNs, occurring perhaps even prior to LN colonization, and highlight their potential for identifying patients at risk for metastatic disease.

## Results

### Combined spatial profiling of proteins and mRNAs in Melanoma sLNs

To construct a spatial atlas of sLNs in melanoma, we assembled a retrospective cohort of 69 melanoma patients who underwent a sLN biopsy as part of their routine clinical diagnosis of the primary tumor. Of these, 39 had negative LNs whereas 30 had metastases in their LNs. Patients were selected for the study such that in both groups roughly half of the patients developed distant metastases within a followup of at least five years (**Fig. 1A**). We named the patient groups using a two-letter code: the first describes whether or not metastases were evident in their sLNs at diagnosis (P-positive/N-negative; P/N), and the second whether the patient proceeded to develop distant metastases within a followup of at least five years (P/N) (**Fig. S1A**). As expected, the tumor’s Breslow score and the patient’s age correlated with recurrence (**Fig. S1B**). A complete summary of the clinical information for all patients is available in **supplementary table 1**.

**Figure 1:**
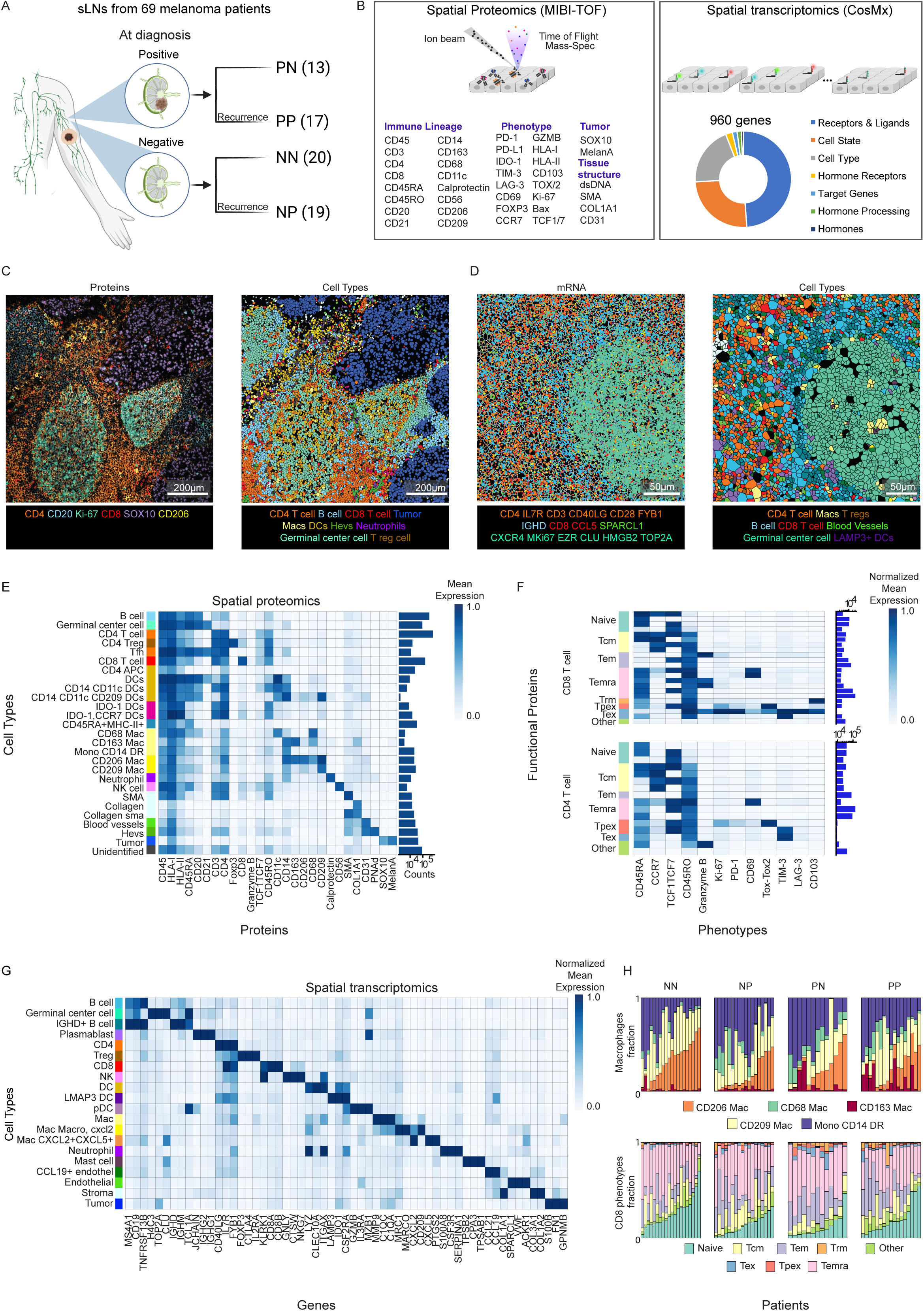
Combined spatial profiling of proteins and mRNAs in Melanoma sLNs. **(A)** The cohort consists of retrospective sLN specimens from 69 melanoma patients. Patients were divided into four groups based on the metastatic status of their nodes (**P**ositive/**N**egative) and whether they developed distant metastases within a followup of at least five years of diagnosis (**P**ositive/**N**egative). **(B)** sLN specimens underwent spatial profiling of 39 proteins using MIBI-TOF (left) and 960 mRNAs using CosMx (right). **(C)** Multiplexed protein images underwent cell segmentation and classification. Shown is a representative image colored by protein expression (left) and cell type classification (right). **(D)** Same as (C) for the mRNA images. **(E)** Heatmap depicts the normalized expression of each protein (x-axis) in each cell type (y-axis) in the protein data. n=68 patients. **(F)** Normalized expression of each protein (x-axis) across CD8 T cell phenotypes (y-axis, top) and CD4 T cell phenotypes (bottom). Phenotypes are grouped together to functional states including Naïve, T central memory (Tcm), T effector memory (Tem), CD45RA+ T effector memory (Temra), Progenitor exhausted T cells (Tpex), Exhausted T cells (Tex), and T resident cells (Trm). **(G)** Heatmap displays the normalized mRNA expression of highly differentially expressed mRNAs (x-axis) across cell types (y-axis). n=33 patients. **(H)** Top: composition of macrophages (y-axis) per patient (x-axis). Bottom: composition of CD8 T cell states (y-axis) per patient (x-axis). Shown are the cell compositions derived from the protein data.

To study the immune composition and organization in the sLNs using multiplexed proteomics imaging, we constructed a panel of 39 antibodies (**Fig. 1B left**). The panel was designed to identify melanoma cells, distinct immune and stromal cell populations as well as various immune cell states. The specificity and sensitivity of all antibodies was validated on control tissues (**Fig. S1C**). We applied this antibody panel to stain formalin fixed paraffin embedded (FFPE) tissue specimens from our cohort and imaged them using MIBI-TOF ^20^. We acquired 2-3 fields of view (FOVs) per patient, resulting in 177 high-dimensional images, each depicting the spatial expression of 39 proteins *in situ* (**Fig. 1C**). We used Mesmer ^22^ to segment individual cells in the images and CellSighter ^23^ to classify the cells to 26 cell types (**Fig. 1E**). As expected, the majority of the cells within the LN are lymphocytes and only patients from the PP and PN groups had tumor cells, as opposed to patients from the NP and NN groups (**Fig. S1D**). We found several types of B cells composing the follicle and the germinal center structures, CD4 and CD8 T cells composing the T-zone accompanied by DCs and macrophages, and CD206^+^ CD209^+^ macrophages, mostly lining the LN sinuses, specifically the medullary sinus^24^.

Next, we clustered the T cell populations based on expression of designated functional proteins. We distinguished distinct CD8 T cell states, including naïve cells which have not yet encountered specific antigen (CD45RA^+^); central memory T cells (Tcm: CD45RO^+^, CCR7^+^), which are antigen-experienced long-lived memory cells that reside in lymphoid tissues; effector memory T cells (Tem: CD45RO^+^, CCR7^-^) and resident memory cells (Trm: CD45RO^+^, CD103^+^), which are usually present in the circulation and non-lymphoid peripheral tissues respectively; exhausted T cells (Tex: PD-1^+^/TOX^+^/TIM- 3^+^/LAG-3^+^), which have been continuously exposed to antigen and precursor-exhausted T cells (Tpex: PD-1^low^ TCF1^+^ TOX^+^), which retain high proliferative capacities, and co-express exhaustion proteins along with self-renewal proteins such as TCF-1^25,26^. We observed similar phenotypes for CD4 T cells, including precursor exhausted CD4 T cells and potentially cytotoxic GZMB^+^ CD4 T cells ^27^ (**Fig. 1F**).

In addition to the protein images, for 33 patients we also performed paired spatial measurements of mRNA using Nanostring’s CosMx platform ^21^, imaging the expression of 960 genes (**Fig. 1B right**).

We used CellPose ^28^ to segment individual cells and InSituType ^29^ to classify the cells into 20 cell types and states (**Fig. 1D,G**). The data acquired using both modalities yielded an overall similar composition and spatial organization for the major cell lineages, including T cells, B cells and tumor cells (**Fig. 1E,G**). The technologies were complimentary, with the protein data providing higher confidence in the phenotype of any particular cell and the mRNA data adding additional cell types, not included in the protein panel such as plasmablasts, mast cells and plasmacytoid DCs (pDCs). Moreover, the mRNA data exposed a more comprehensive phenotypic landscape within the macrophage and DC populations. For instance, while both the protein and mRNA data identified CD206^+^ macrophages (expressing the MRC1 gene), the latter further revealed that they also express MARCO, CXCL2 and CXCL5 (**Fig. 1. E, G**). Comparison of the cell type composition among the different groups in the cohort revealed distinct differences in macrophages and T cells phenotype, with higher abundance of sinus macrophages (CD206^+^ macrophages, **Fig. 1H top**) in NN patients and increased Naïve CD8 T cell in NP patients. (**Fig. 1H bottom, Fig. S1E**). Altogether, the combination of protein and transcript expression provides a superior identification approach to immune cell types.

To identify microenvironments in the sLNs, we developed a novel deep-learning self-supervised-based approach to cluster patches in the images according to their cellular composition and protein expression. Briefly, images depicting the outputs of the cell classification were fed into DINO ^30^, a self-supervised vision transformer, which clusters together images with similar semantic features (**Fig. 2A**). The model passes two different random transformations of an input image to a student and teacher network. It was trained to position these transformations in proximity in the final embedding space, which we then clustered using PhenoGraph^31^ to produce the final classifications of cell niches.

**Figure 2:**
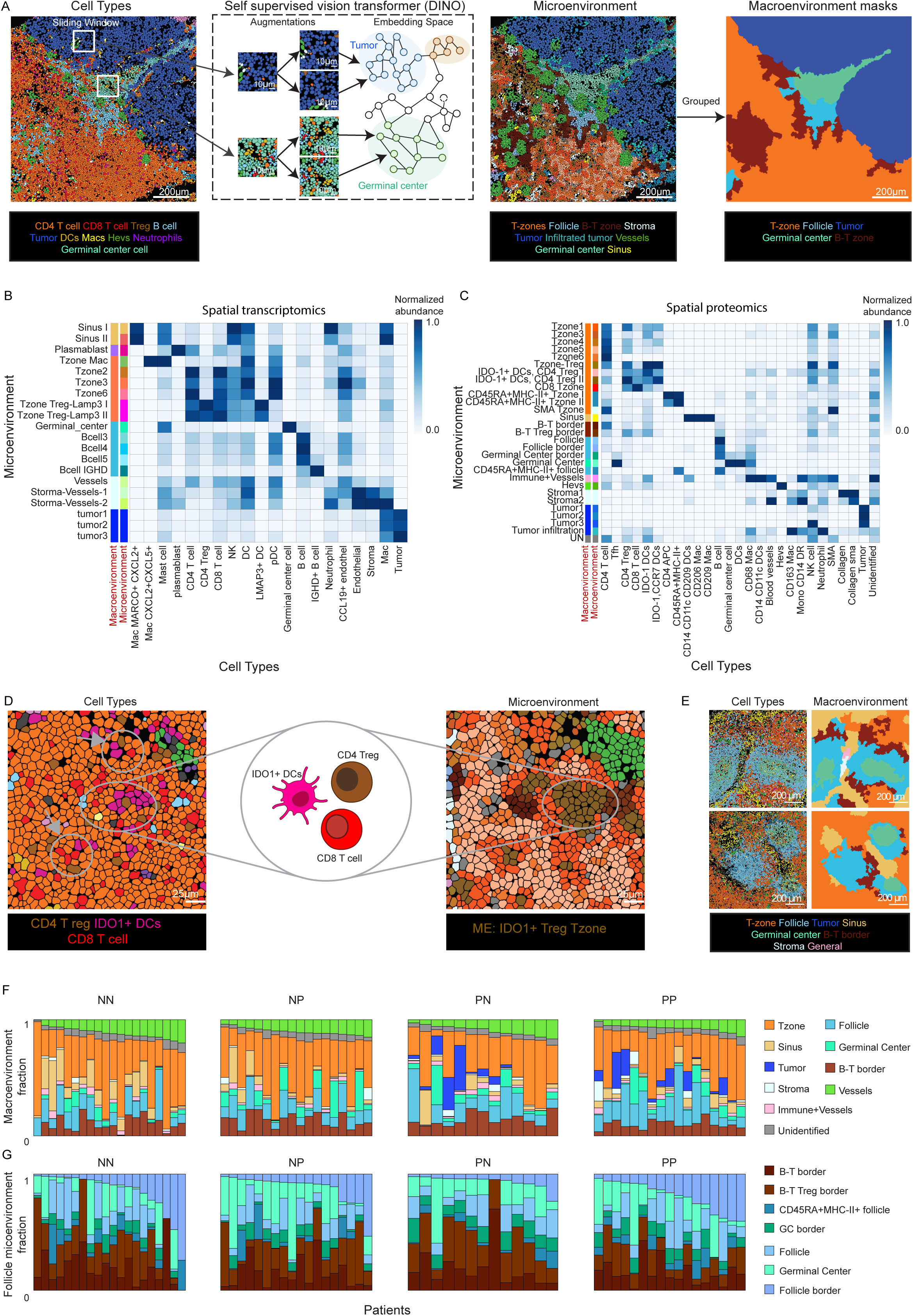
Vision transformer identifies micro and macroenvironments in sLNs. **(A)** Images of classified cells were fed into DINO, a self-supervised vision transformer. DINO was trained by performing augmentations on images and learning to project them in proximity in the embeddings space. After training on sliding windows, the embedding space was clustered into microenvironments. Microenvironments with similar cell type compositions were manually grouped to macroenvironments, which were segmented to generate macroenvironment masks. **(B)** Heatmap depicts the normalized composition of cell types (x-axis) in each microenvironment (y-axis) for the mRNA data. Rows are colored by the macroenvironment (first column) and the microenvironment (second column). **(C)** Same as (B) for the protein images. **(D)** Example image colored by cell type classification (left) and microenvironment classification (right). Circle and arrows emphasize a microenvironment (brown) colocalizing IDO1+ DCs (pink) with Tregs (brown) and CD8 T cell (red). **(E)** Example images colored by cell types (left) and macroenvironments (right), presenting the LN compartments. The cell type colormap matches that of 1E. **(F)** Composition of macroenvironments (y-axis) per patient (x-axis), derived from the protein data. **(G)** Composition of the microenvironments comprising the follicle macroenvironment (y-axis) per patient (x-axis), derived from the protein data.

We identified 20 microenvironments in the mRNA data and 28 microenvironments in the protein data (**Fig 2. B,C**). We annotated each microenvironment according to its main cell types. Although not all cell types were identical between the mRNA and protein data, we could identify recurring motifs. For example, the sinuses were enriched for CD206+ macrophages in the protein data and CD206^+^ CD209^+^ MARCO^+^ CXCL2^+^ macrophages in the mRNA data. Both data types also identified a microenvironment enriched for interactions of DCs and Tregs, defined by colocalization of LAMP3^+^ DCs with Tregs in the mRNA data and an equivalent microenvironment co-localizing IDO-1^+^ DCs and Tregs in the protein data (**Fig. 2D**). We then manually grouped microenvironments into macroenvironments describing typical regions of the LN such as the B cell follicles, interfollicular T cell zones (T-zone) and the LN’s sinus (**Fig. 2E, 2F, S2A**). We further distinguished different layers of the B cell follicles, including the germinal center and mantle zone, which was enriched for naïve B cells. The border of the follicle was defined by a macroenvironment enriched by a mixture of B cells and T cells (B-T border) (**Fig. 2E**). The T-zone was also composed of several micro-environments characterized by local enrichments of CD4 T cells, CD8 T cells, Tregs, DCs and APCs (**Fig. 2B,C**).

Comparing the microenvironment and macroenvironments composition between the different groups revealed a higher fraction of the sinus macroenvironment in NN patients (**Fig. 2F**). In the follicles, we found an enrichment of germinal center structures in NP patients compared to NN, and an increased abundance of Treg-enriched B-T border in PN patients compared to PP patients (**Fig. 2G**). Overall, our multiplexed proteomics and transcriptomics datasets allowed us to extract features on cell types, cell states and cell niches in sLNs.

### Tumor seeding associates with immune remodeling in the lymph node

To understand the microenvironment of melanoma metastases in the sLN, we first focused on patients who had positive sLNs (PN and PP, subsequentially referred to as PX). We analyzed the metastatic regions and compared the composition and phenotype of non-tumor cells inside and outside of the metastases (**Fig. 3A**). This analysis revealed a unique intra-metastasis microenvironment, associated with a preferential immune infiltration of macrophages, neutrophils and CD8 T cells (p<0.05, **Fig. 3B,C)**. This immune composition was very different than the LN composition outside the metastases, which was enriched for CD4 T cells and B cells. We also observed an increase of stromal cells inside the metastasis and lack of the high endothelial venules (HEVs) characteristic of the lymph node parenchyma, accompanied by an increase in metastasis-associated blood vessels (p<0.001, **Fig. 3C,D, Fig. S2B**). These results suggest that although LN metastases are seeded in immune hubs, they drive a distinct local immune microenvironment.

**Figure 3:**
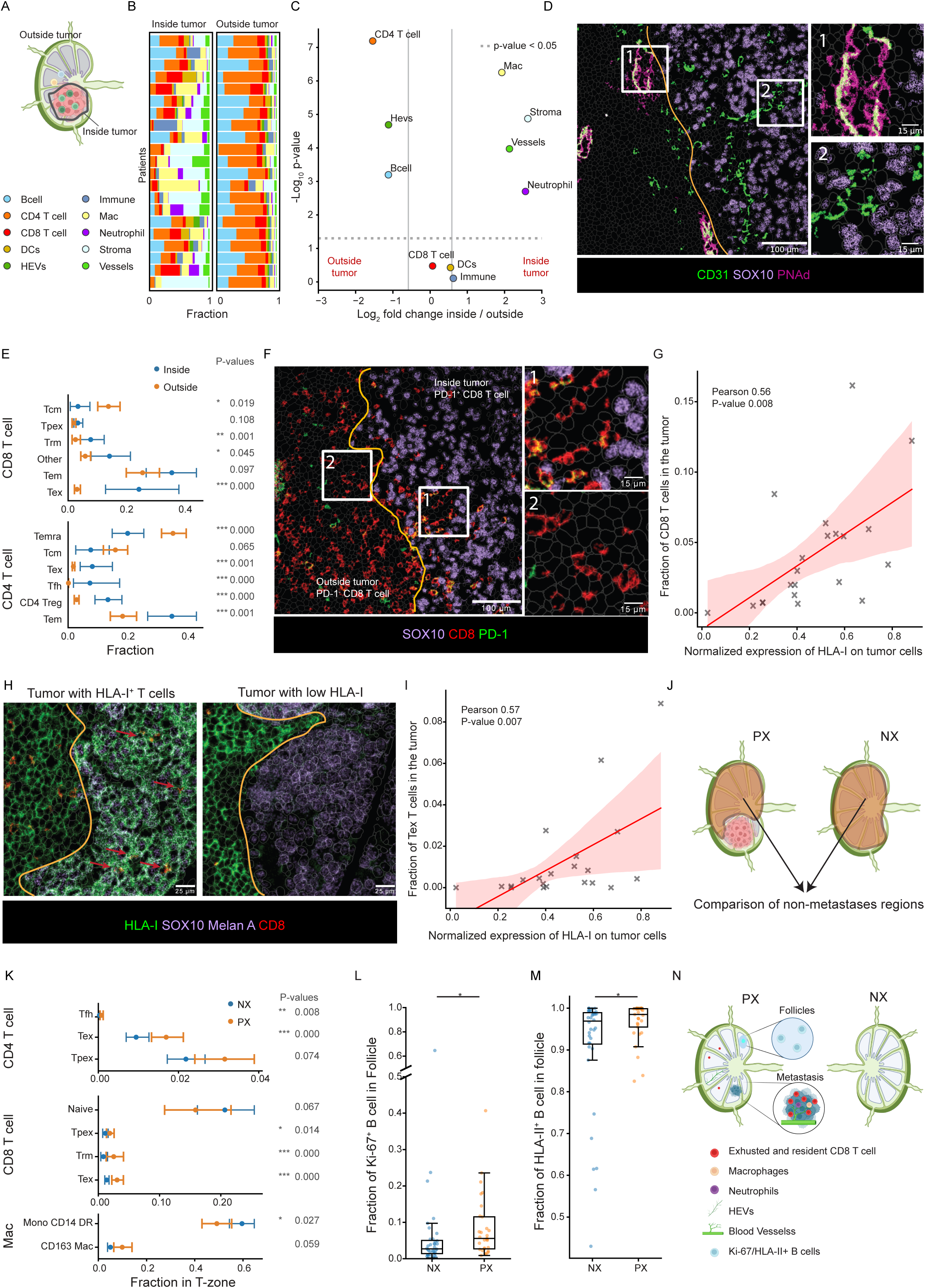
Tumor seeding associates with immune remodeling of the lymph node. **(A)** Illustration, inside and outside the metastatic region. **(B)** Cell type composition (x-axis) per patient (y-axis) inside (left) and outside (right) the metastatic macroenvironment. n=21 patients. **(C)** Volcano plot of cell type abundances inside versus outside the metastatic region. n=21 patients. **(D)** Example image depicting blood vessels (CD31, green) inside the metastases and HEVs (CD31 and PNAd) outside. **(E)** Comparison of CD8 and CD4 T cell phenotypes (y-axis) within the metastases versus the T-zone. The X-axis shows the fraction of cells out of their respective lineage. P-values were calculated across patients. Lines represent the 95% confidence intervals, and dots indicate the mean values across patients. Only patients with least 10 CD4 or CD8 T cells were included in the analysis. CD8 T cell analysis: 17 patients inside, 30 outside. CD4 T cell analysis: 20 patients inside, 30 outside. **(F)** Representative image of Tex CD8 T cells (CD8+ PD-1+) within the tumor. **(G)** Correlation between the normalized mean expression of HLA-I+ on tumor cells (x-axis) and CD8 T cell infiltration inside the tumor macroenvironment (y-axis). n=21 patients. **(H)** Representative images of an HLA-I+ tumor with CD8 infiltration (left) vs an HLA-I low Tumor without CD8 T cell infiltration (right). **(I)** Correlation between the normalized mean expression of HLA-I+ on tumor cells (x-axis) and the fraction of Tex T cell infiltration within the tumor macroenvironment (y-axis). n=21 patients. **(J)** Illustration of non-metastases regions in a clean (NX) vs metastatic (PX) lymph node. **(K)** Same as (E), comparing CD4 T cell, CD8 T cell and macrophage phenotypes within the T-zone macroenvironment of NX (blue) and PX (orange) patients. PX, NX=30, 38 patients. **(L)** Fraction of Ki-67+ B cells out of all B cells in the follicles of PX and NX patients, calculated for each patient with a minimum of 10 B cells. Each dot represents an individual patient. PX, NX=29, 36 patients. **(M)** same as (L) for HLA-II+ B cells. PX, NX=29, 36 patients. **(N)** Illustration summarizing the key characteristics of NX vs PX. **All p-values were calculated using the Mann-Whitney statistical test. The analysis was limited to regions with more than 100 cells.**

Next, we focused on the phenotypes of CD8 and CD4 T cells inside and outside the metastasis regions. For each patient we calculated the fraction of each distinct immune phenotype out of the cells of the same type. For example, we calculated the fraction of exhausted CD8 T cells out of the CD8 T cells, either inside or outside the metastases microenvironments. Across patients, we observed more effector (p=0.001), exhausted (p=0.001) and T regs (p<0.001) out of all the CD4 T cells inside the metastases (**Fig 3E**). Similarly, for CD8 T cells we observed more resident (p=0.001) and exhausted phenotypes (p<0.001) inside the metastasis. This difference diminishes when we compare the precursor exhausted (Tpex) population (p=0.11), suggesting that the progression of exhaustion occurs in the context of the metastasis. Interestingly, for some patients we observe a sharp boundary, in which T cells within the metastasis express exhaustion proteins, such as PD-1, whereas T cells outside do not (**Fig. 3F**). This behavior is accompanied by PD-L1 expression in the metastasis microenvironment (**Fig. S2C**). Together, these observations suggest that exhaustion occurs within the metastatic niche. The paucity of exhausted cells outside the metastatic niche may also suggest that the exhausted T cells remain within the metastasis and there may be little efflux of T cells from the metastasis microenvironment to the metastasis-adjacent regions.

Our analysis of CD8 T cell infiltration into the metastases, revealed variability across patients, spanning over a 15-fold difference in abundance (**Fig. 3B, G**). To explore potential mechanisms underlying this variance, we compared the degree of CD8 T cell infiltration to the phenotypes of tumor cells. We found that expression of HLA-I on the tumor cells was positively correlated with CD8 T cell infiltration into the metastasis (R=0.56 p=0.008, **Fig. 3G, H**). Furthermore, we found that HLA-I expression on tumor cells was correlated with an exhausted phenotype on the infiltrated CD8 T cells (R=0.57 p=0.007, **Fig. 3I**). These results suggest that continuous stimulation by antigens in the metastasis microenvironment drives T cell exhaustion, and that downregulation of HLA-I by tumor cells is an effective mechanism to evade predation by CD8 T cells, as previously suggested^32,33^.

Our results thus far have shown that metastases remodel their immediate surroundings. Next, we evaluated whether the presence of metastases in the LN correlates with immune remodeling outside the metastases. To this end, we compared the non-metastatic regions of the sLN between patients who had metastases in their LN (noted by PX) to patients who did not have metastases in their LNs (noted by NX) (**Fig. 3J**). When comparing the distribution of phenotypes of cells in the T-zone, we observed an increase in the populations of resident T cells (CD8 T cell p<0.001) and exhausted T cells (CD8 T cell p<0.001) in patients with metastases in their LNs (**Fig. 3K**), indicating higher activation of CD4 and CD8 T cells in metastatic lymph nodes. Comparing the phenotypes of macrophages revealed a more naïve-like population of monocytes expressing HLA class II (Mono CD14 DR; p=0.027) in the negative LNs (**Fig. 3K**).

Next, we compared the follicle macroenvironment. We observed an increase of Ki-67 on B cells in the follicle (p=0.01, **Fig. 3L**), suggesting more activated germinal centers in LNs that harbor metastases. Accordingly, we found that metastatic sLNs had slightly larger follicles compared to negative sLNs (ns, **Fig. S2D**). We also found altered phenotypes in B cells in metastatic LNs, which had a higher fraction of cells expressing HLA class II (p=0.03, **Fig. 3M**), and a higher fraction of B cells expressing CD11c (p=0.012, **Fig. S2E**). The latter subset may represent atypical B cells, previously connected to chronic infection, systemic autoimmunity and memory formation^34–37^. In contrast, negative LNs had more suppressive PD-L1^+^ DCs (p=0.036, **Fig. S2F**). Overall, metastatic LNs showed signs of immune remodeling, suggesting that the LN is globally shaped by the presence of the metastases.

### Immune organization in patients with LN metastases is predictive of development of distance metastases

Since we observed that the presence of metastases remodels the LN both locally and globally, we proceeded to analyze these groups separately to assess if immune responses in the LN associate with future development of distant metastases. First, we focused on the patients who had metastatic sLNs, and compared the immune landscape between the patients who proceeded to either develop distant metastases (PP) or not (PN) (**Fig. 4A**). While the cell lineage composition was similar in PN and PP patients (**Fig. S3A**), analysis of the phenotypes of the CD8 T cells in the metastatic regions revealed that in PN patients they are characterized by a more stem-like memory profile co-expressing TCF, CD45RO and granzyme B (PN=2.7%, PP=0.7%, p=0.024, **Fig. 4B, C**). Accordingly, the metastatic regions of PN patients had more CD103^+^ (PN=13%, PP=2.2%, p=0.016) and TCF^+^ DCs (PN=15%, PP=4%, p=0.014), which could serve as local drivers of T cell activation^10,11^ (**Fig. 4D**). In contrast, we found more M2-like macrophages expressing CD209 (PN=16%, PP=34%, p=0.039) and CD163 (PN=23%, PP=44%, p=0.06) in the metastatic regions of PP patients, previously shown to have an immunosuppressive effect^38,39^ (**Fig. 4D**). Altogether, in the metastatic regions of PN patients we observed activation of T cells, possibly by local DCs, in contrast to potentially suppressive macrophages in PP patients.

**Figure 4:**
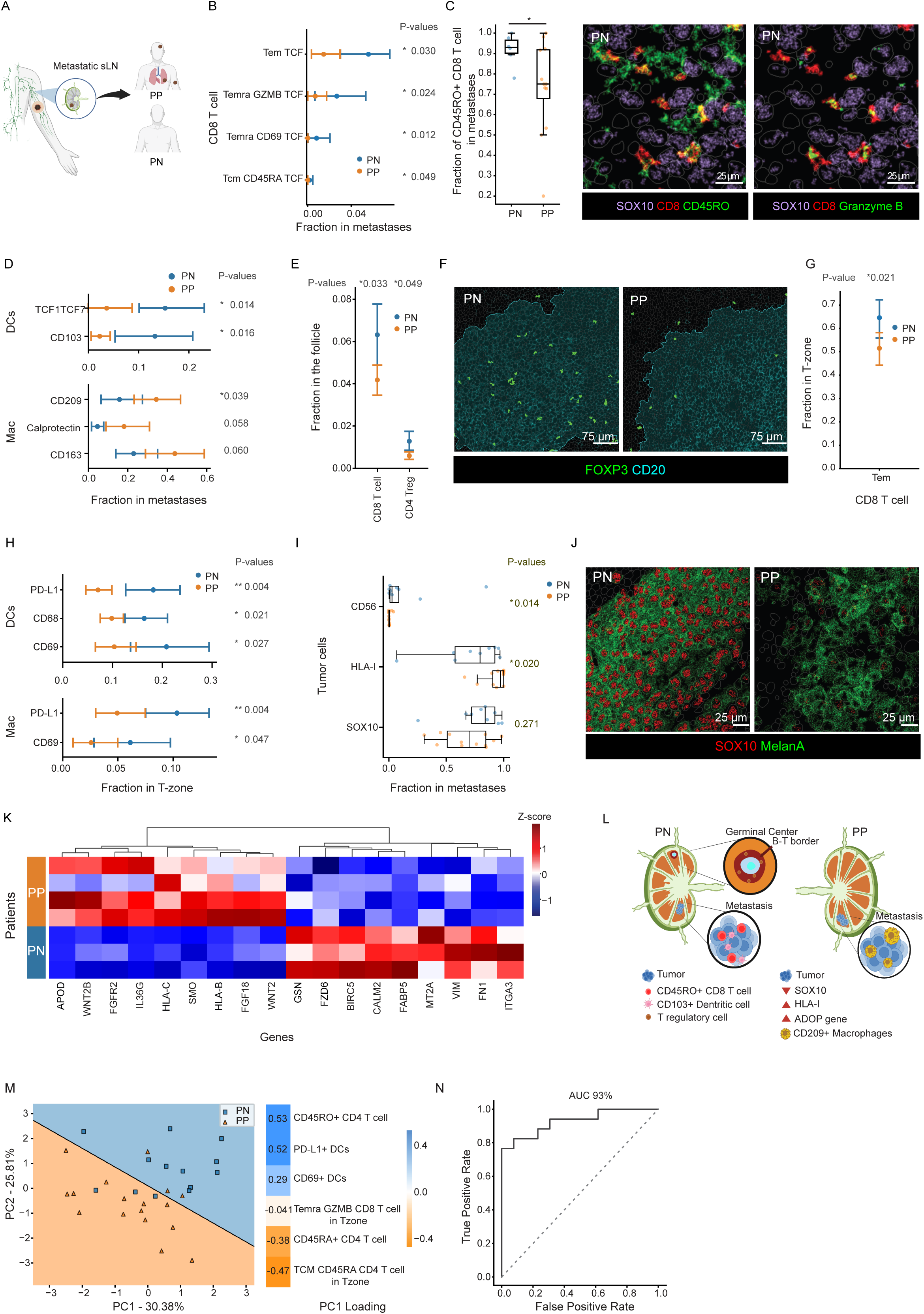
Immune organization in patients with LN metastases is predictive of development of distant metastases. **(A)** Illustration of metastatic lymph nodes; PP patients developed distant metastases while PN did not. **(B)** Comparison of CD8 T cell phenotypes (y-axis) within the metastases between PN (blue) and PP (orange) patients. The X-axis shows the fraction of cells out of their respective lineage. P-values were calculated across patients. Lines represent the 95% confidence intervals, and dots indicate the mean values across patients. Only patients with at least 10 CD8 T cells were included in the analysis. PN, PP=6, 12 patients. **(C)** Left: fraction of CD45RO+ CD8 T cells out of all CD8 T cells in the metastatic regions of PN vs PP patients, calculated for each patient with a minimum of 10 CD8 T cells. Each dot represents a patient. PN, PP=6, 12 patients. Right: example protein images showing CD8 T cells (red) expressing CD45RO (green, left) or Granzyme B (green, right) next to tumor cells (SOX10; light purple). **(D)** Same as (B), showing a comparison of positivity fractions (x-axis) for different proteins (y-axis) in DCs and Macrophages within the metastases. Calculated for each patient with a minimum of 5 DCs/Macrophages. DCs analysis: PN, PP=6,8 patients. Macrophages analysis: PN, PP=9,12 patients. **(E)** Same as (B), showing the fraction of CD8 T cell and T regulatory cells in the follicles. PN, PP=13, 17 patients. **(F)** Representative images of follicles in PN (left) and PP (right). FOXP3 signal (green) is dilated for emphasis. The follicle macroenvironment is outlined. **(G)** Same as (B) in the T-zone macroenvironment. The Tem and Temra CD8 phenotypes were grouped in this analysis. PN, PP=13, 17 patients. **(H)** Same as (D) in the T-zone macroenvironment. PN, PP=13, 17 patients. **(I)** Same as (D), comparing the protein positivity fraction in tumor cells within the metastases of PN and PP patients. PN, PP=9, 12 patients. Only patients with a minimum of 10 tumor cells were included in the analysis. **(J)** Representative images showing a high SOX10 PN tumor (left) and a low SOX10 PP tumor (right). **(K)** Heatmap depicting the normalized mRNA expression of tumor cells per patient (y-axis). The X-axis displays the top differentially expressed genes between PN and PP. Only regions with at least 20 tumor cells were included. PN, PP=3, 4 patients. **(L)** Illustration summarizing the key characteristics of PN and PP. **(M)** Left: PCA of PN (blue squares) and PP (orange triangles) patients based on six selected features. The X and Y axes show the first and second principal components, respectively. The separation line was determined using a support vector machine (SVM). Right: PCA loadings for the first component. **(N)** ROC curve of a leave-one-out analysis over all PX patients, predicting whether they will develop distant metastases. PN, PP=13, 17 patients. **All p-values were calculated using Mann-Whitney statistic test. For (B-D, E-I) the analysis was limited to regions with more than 100 cells.**

Subsequently, we analyzed the cellular composition outside the metastatic regions to identify global differences in the sLNs of PN and PP patients. Comparing the follicle regions, we identified more blood vessels in PP patients (p=0.01, **Fig. S3B**) and increases in Tregs (PN=1.2%, PP=0.6, p=0.049) and CD8 T cells (PN=6.3%, PP=4.2%, p=0.033) in the follicles of PN patients (**Fig 4 E,F**). It was also manifested by an increase in the microenvironment that colocalized Tregs, CD8 T cells and IDO-1 DCs in the B-T cell border of the follicle of PN patients (p=0.01, **Fig. S3C**). Next, we compared the T-zones, and in accordance with the results in the metastases, we observed that PN patients had more effector memory CD8 T cells (PN=65%, PP=51%, p=0.021, **Fig. 4G**). We also found higher expression of PD-L1 (PN=18%, PP=7%, p=0.004) and CD69 (PN=21%, PP=10%, p=0.027) in DCs and macrophages in PN patients (**Fig. 4H**). Those PD-L1 expressing cells might participate in driving the exhaustion of the CD8 T cells^40^. Overall, analysis of immune phenotypes in both metastatic and non-colonized LN compartments revealed distinct states: patients who developed distant metastases showed suppressive myeloid cells, while those who remained metastasis-free exhibited T cell responses, potentially impaired by T regulatory cells.

Next, we explored properties of the tumor cells and associated them with the observed immune organization and development of distant metastases. In the protein data, PN metastases had moderately increased expression of CD56 (p=0.014, **Fig. 4I**), which may indicate neuroendocrine differentiation^41^. They also exhibited reduced expression of HLA-I, potentially as a mechanism of acquired resistance to the T cell responses^42^ (p=0.02, **Fig. 4I**). A subset of PP patients showed reduced expression of SOX10, a transcription factor driving neural crest differentiation, supporting previous results that loss of SOX10 associates with a more invasive phenotype^43^ (**Fig. 4I,J**). Similarly, analysis of the mRNA data in a subset of patients confirmed higher expression of HLA-I in tumor cells in PP patients. In addition, PP patients exhibited higher expression of APOD and SMO, which were previously linked to invasion ^44^ ^45^. Moreover, PP patients showed elevated expression of fibroblast growth factors and their receptor. Melanoma has been shown to stimulate FGFR in an autocrine manner, proposed as a self-sustaining mechanism that promotes tumor survival and proliferation ^46^ (**Fig. 4K)**. Tumors from PN patients displayed increased expression of BIRC5, CALM2 and FABP5, all of which have been linked to tumor proliferation, the inhibition of apoptosis, or both ^47–51^. Additionally, these tumors exhibited increased expression of FN1 and ITGA3, which are associated with migration^52,53^ (**Fig. 4K**). Altogether, analysis of tumor phenotypes revealed downregulation of antigen presentation, perhaps as a defense against T cell predation, but it did not clarify why T cell responses were less prominent in PP patients.

Clinically, systemic treatment is guided by the Breslow thickness of the primary tumor, the metastasis size in the LN and relevant clinical factors related to risks of toxicity. Our analysis identified features in metastatic sLNs that correlate with future development of distant metastases **(Fig. 4L)**. We therefore assessed the predictive power of multiplexed imaging for forecasting recurrence. We selected six features that were independently predictive of metastatic recurrence, based on phenotypes, macroenvironments and protein expression, and which could be obtained using low-plex imaging. We then preformed principal component analysis (PCA) to extract correlated components that describe the data (**Fig. 4M).** We then repeated this pipeline, each time leaving out one of the patients. In each iteration we performed principal component analysis and trained a Support Vector Machine (SVM) classifier with the first two PCs as features. We proceeded to test our classifier on the held-out patient. Overall, our approach resulted in an AUC of 93% (**Fig. 4N**), compared with an AUC of 56% when using age and Breslow thickness (**Fig. S3E**). In our cohort, around 40% of the PP patients were not treated with systemic therapy, such as immune checkpoint inhibitors (**Fig. S3D**). Using our classifier, 88% of them would be classified as likely to develop distant metastases. Our results suggest that multiplexed imaging of the sLNs has the potential to be used to identify patients with a high chance of developing distant metastases.

### Immune organization in patients with negative sLNs is predictive of development of distant metastases

Next, we focused on patients with negative lymph nodes, who pose a challenging treatment dilemma. While some benefit from immune checkpoint blockade (ICB)^8^, many do not require treatment, risking unnecessary morbidity, mortality, toxicity, and significant financial costs. We compared the patients who proceeded to either develop distant metastases (NP) or not (NN) (**Fig. 5A**), and analyzed the composition of microenvironments as well as the composition of cell types and phenotypes in each macroenvironment. We found that overall NP patients were characterized by a T-cell mediated immune response in the LN compared to NN patients. NN patients displayed more naïve T cell phenotypes, for both CD4 and CD8 T cells (p<0.05, **Fig. 5B**). In contrast, NP patients had more central memory CD8 and CD4 T cell phenotypes (p<0.05, **Fig. 5B)**, overall suggesting the presence of antigen-experienced circulating T cells. Accordingly, we observed increased CCR7 expression in the T-zone for both CD4 T cells (p=0.042) and DCs (p=0.013) in NP patients (**Fig. 5C**), which promotes the migration and homing of immune cells to lymphoid tissues. In addition, LNs from NP patients had several indications for immunosuppression, such as more T regulatory cells (Tregs), which have been shown to have a suppressive effect on the immune response in the LNs ^54^ (p=0.042, **Fig. 5B)**, and more DCs expressing PD-L1, known to induce a suppressive microenvironment ^55^, in the T-zone (p=0.056, **Fig. 5D).** Collectively, these results could suggest that either suppressive DCs from the primary tumor are seeding the LN and inducing a tolerogenic immune environment in the T zone or that sLNs of NP patients were predisposed to tolerance (**Fig. 5N**). The mRNA data also indicated increased IDO-1 expression in NP patients, further supporting the enrichment of suppressive DCs in this group (**Fig. S4A**). In contrast, DCs in NN patients exhibited higher levels of TIMP1. Increased TIMP1 expression in the immune compartments of the tumor microenvironment in melanoma patients was recently associate with good prognosis^56^.

**Figure 5:**
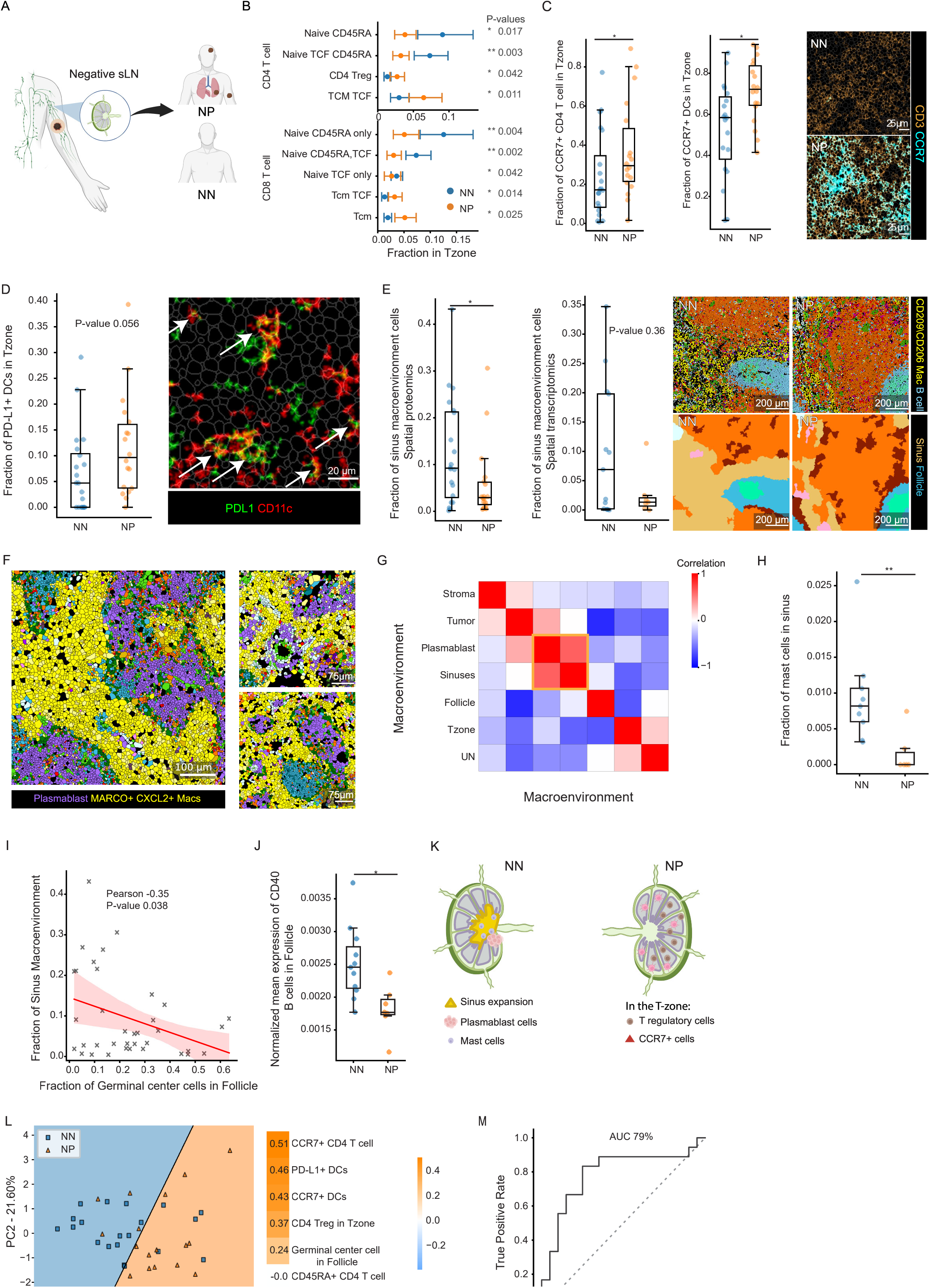
Immune organization in patients with negative sLNs is predictive of development of distant metastases. **(A)** Illustration of a negative lymph node; NP patients developed distant metastases while NN did not. **(B)** Comparison of CD4 and CD8 T cell phenotypes (y-axis) within the T-zone between NN (blue) and NP (orange) patients. The X-axis shows the fraction of cells out of their respective lineage. P-values were calculated across patients. Lines represent the 95% confidence intervals, and dots indicate the mean values across patients. Only patients with at least 10 CD4/CD8 T cells were included in the analysis. NN, NP=20, 18 patients. **(C)** Left: fraction of CCR7+ cells out of the CD4 T cells in the T-zone of NN and NP patients. Only patients with a minimum of 10 CD4 T cells were included in the analysis. NN, NP=20, 18 patients. Middle: same for DCs. Right: representative images of CCR7+ T cells in the T-zone of NN and NP patients. NN, NP=19, 18 patients. **(D)** Left: fraction of PD-L1+ DCs out of all DCs in the T-zone of NN and NP patients. Each dot represents an individual patient. Only patients with a minimum of 5 DCs were included in the analysis. NN, NP=20, 18 patients. Right: Representative image of PD-L1+ DCs in the T-zone of an NP patient. **(E)** Fraction of the sinus macroenvironment cells in NN and NP patients, with each dot representing a patient. Left: Spatial proteomics. Middle: Spatial transcriptomics. Right: Representative images showing a larger sinus macroenvironment in NN patients. Images are colored by cell types (top row) or macroenvironments (bottom row). Cell types and macroenvironments are colored as in figures 1C and 2A respectively. For spatial proteomics analysis: NN, NP=20,18 patients, for spatial transcriptomics analysis: NN,NP=13,7 patients. **(F)** Representative images showing proximity between sinus macrophages and plasmablast cells. Cell types are colored as in figures 1G. **(G)** Heatmap showing the correlation of macroenvironments across fields of view. The yellow box highlights the correlation between the sinuses and plasmablasts macroenvironments. n=77 fovs from 33 patients. **(H)** Fraction of Mast cells in the sinusoidal regions of NN and NP patients. Each dot represents an individual patient. NN,NP=9,6 patients. **(I)** Anti-correlation between the fraction of germinal center cells in the Follicle (x-axis) and fraction of sinus macroenvironment (y-axis). Each dot represents an individual patient. n=36 patients. **(J)** Fraction of CD40+ B cells out of the total B cell population within the follicle macroenvironment. Only patients with a minimum of 10 cells were included in the analysis. NN, NP=11,7 patients. **(K)** Illustration summarizing the key characteristics of NN and NP. **(L)** Left: PCA of NN (blue squares) and NP (orange triangles) patients based on 7 selected features. The X and Y axes show the first and second principal components, respectively. The separation line was determined using a support vector machine (SVM). Right: PCA loadings for the first component. **(M)** ROC curve of a leave-one-out analysis over all NX patients, predicting whether they will develop distant metastases. NN, NP=20, 18 patients. **All p-values were calculated using Mann-Whitney statistic test. For (B-D,H-I) the analysis was limited to regions with more than 100 cells.**

In contrast to NP patients, NN patients had an increase in the macroenvironment of the sinusoids (p=0.045, **Fig. 5E**), also manifested by a global increase in CD206^+^ macrophages **(Fig. 1H)**. This expansion was correlated with an increase in the abundance of blood vessels in the T zone (R=0.79 p<0.001, **Fig S4B**). Together, these results suggest that the sLNs of NN patients are preferentially undergoing an expansion of the sinusoids and lymphangiogenesis, which have been shown to follow an immune response to inflammation and accompany the egress of lymphocytes from the LN^57^. Accordingly, in the mRNA images, we observed a high correlation between the expansion of the sinusoids and a macroenvironment that was enriched with plasmablasts (R=0.76, p<0.01, **Fig 5F,G**, **Fig. S4C)**. These plasmablasts were located in close proximity to the sinusoids (**Fig. 5F,G**), possibly indicating their imminent exit from the LN. The sinusoids in NN patients were also enriched for mast cells (**Fig. 5H**), which have previously been associated with good prognosis, in both primary tumors and LN metastases^58,59^. Taken together, these findings indicate an immune response in NN patients that involves an expansion of the sinusoids and likely the egress of plasmablasts from the LN.

The spatial organization of the LNs in NN patients was reminiscent of an extrafollicular response (EFR), in which plasmablasts are produced from the extrafollicular zone with or without a limited germinal center response, with limited T cell dependent activation of B cells^60^. In support of this hypothesis, we found a negative correlation between the expansion of the LN’s sinus and the abundance of follicular germinal center cells in the follicle (R=-0.35, p=0.038, **Fig. 5I)**. The higher abundance of mast cells in the sinuses of NN patients (**Fig. 5H**) further supports this hypothesis as mast cells have been demonstrated to enhance the differentiation of naïve B cells into plasmablasts^61^. Conversely, we observed an increased presence of CD40+ B cells in the follicle of NN patients (p=0.02, **Fig. 5J**) in the mRNA data, which could suggest that there is a T cell-dependent stimulation of B cells in the follicles of NN patients^62^.

To conclude, we suggest that the lymph nodes of NP patients exhibit increased influx of DCs and T cells, whereas NN patients show expansion of the sinusoids and may be enriched for extrafollicular responses **(Fig. 5K)**. We note that in accordance with previous work^63^ in this cohort the average age of NP patients (63 y/o) was higher than the average age of NN patients (53.8 y/o) (**Fig. S1B left**). As such, age may be associated with some of the differences between NP and NN. For example, previous studies have shown an increase in the accumulation of Tregs in the LNs with age ^64^, and we also found a correlation between age and accumulation of Tregs in the LN (R=0.38, p=0.017, **Fig. S4D**). In contrast, expansion of the sinusoids was not correlated with patient age (R=0.03, p=0.837, **Fig. S4E**), suggesting it as an age-independent differentiating factor between NN and NP patients.

Finally, we examined whether the spatial changes between NN and NP patients could predict whether a patient with negative sLNs will develop distant metastases. For this analysis we focused on the MIBI data, for which we had more samples. We selected 7 features that were independently predictive of metastatic recurrence, and could be obtained using low-plex imaging, and preformed PCA to extract correlated components that describe the data (**Fig. 5L).** The first component is mostly comprised of CCR7^+^ CD4 T cells and the expansion of the sinusoids. We then repeated this pipeline, each time leaving out one of the patients. In each iteration we performed feature selection, PCA and trained a Support Vector Machine (SVM) classifier. We then tested our classifier on the held-out patient. Overall, our approach resulted in an area under the curve (AUC) of 79% **(Fig. 5M)** compared with an AUC of 74% when using age and Breslow depth **(Fig. S4F)**. Under standard of care, patients with negative LNs are generally not treated with systemic therapy, such as immune checkpoint inhibitors, but recent clinical trials have indicated that stage IIb/c melanoma patients could benefit from immunotherapy^8^. Our results suggest that in depth analysis of the sLNs could be used to identify patients with a high chance of developing distant metastases, who could potentially benefit from systemic therapy.

## Discussion

We analyzed the spatial organization of sentinel lymph nodes in melanoma patients using multiplexed protein and RNA imaging techniques. Combining data from both methods proved complementary. Multiplexed proteomics provided high-quality data and allowed to profile more samples and to reliably assess the immune cells phenotypes but was limited by the number of proteins that could be analyzed. Conversely, spatial transcriptomics offered richer molecular insights but reduced sample throughput and sacrificed the quality of individual cell signals due to low mRNA counts. Together these methods were synergistic.

Our findings revealed that within lymph nodes, melanoma metastases create a unique immune microenvironment enriched with myeloid cells, neutrophils and exhausted cytotoxic T cells. Notably, precursor-exhausted and exhausted T cells were primarily confined to metastatic niches, suggesting a suppressive local environment that triggers rapid T cell exhaustion. This observation aligns with studies in mice showing that CD8+ T cell dysfunction is established within hours of encountering tumor antigens ^65^. Alternatively, it could indicate limited movement of T cells from metastatic to adjacent regions, supported by previous research showing that T cell motility inversely correlates with tumor density ^66^. Future studies tracking individual T cells in the tumor microenvironment are needed to clarify these possibilities.

Our analysis identified distinct immune architectures among different patient groups, which were associated with the development of distant metastases. In patients with sLN metastases, we found an inverse correlation between activated CD8 T cells in metastatic regions and the development of distant metastases. Since approximately 50% of this group were subsequently treated with immunotherapy, our findings align with studies showing that pre-existing PD-1^+^ CD8 T cells in the metastases are associated with positive therapeutic outcomes and survival^67^. Furthermore, PN patients showed an increase in CD103^+^ DCs in their metastases, potentially driving T cell activation^10,11^. Their metastases exhibited reduced expression of HLA-I, possibly to evade T cell mediated killing. Together these observations point to an anti-tumor T cell response. Perhaps paradoxically, PN patients also had more Tregs in the follicle regions. These Tregs could be T follicular regulatory cells, which are known to reside on the border of the follicle and could regulate the germinal center response ^68^. It could indicate that at the time of biopsy, the anti-tumor T cell response has become tolerogenic. Immune checkpoint blockade, given to 50% of PN patients in our cohort, may help reverse this process.

In contrast, patients with LN metastases who developed distant metastases had weaker signs of T cell activation in the metastasis regions. Instead, we observed an increase in CD163^+^ and CD209^+^ macrophages, reinforcing the connection between M2-like macrophages in metastases and poor prognosis^38,39^. A subset of these patients also showed reduced expression of SOX10, suggesting a less differentiated tumor state. Together these findings raise the hypothesis that these aggressive metastases instigate weaker anti-tumor T cell responses, although our protein and mRNA panels lacked sufficient targets to determine the underlying mechanisms. Future research should explore whether immune evasion is driven by a lack of neoantigens or other suppressive pathways in the tumor or stroma.

Our study also identified features predictive of future development of distant metastases in patients with negative LNs. Patients that developed distant metastases had increased expression of CCR7 and more Tregs, potentially promoting tolerance. In our cohort NP patients tended to be older than NN patients. Age is a known poor prognostic factor, and is also linked to Treg accumulation in secondary lymphoid organs ^64,69^. Our results suggest that Tregs may be a link between age and worse prognosis. Interestingly, whereas a T cell response was positively associated with development of distant metastases in patients with negative lymph nodes (NP patients), it was negatively associated with development of distant metastases in patients with metastatic lymph nodes (PN patients). This discrepancy may partially stem from differences in treatments between these groups, making them difficult to compare: NP patients were not treated with immunotherapy, whereas ±50% of PN patients were treated and ±75% of PN patients had axillary lymph node dissection. This could suggest that patients with negative lymph nodes that exhibit signs of suppressive T cell responses could benefit from immunotherapy. Indeed, recent clinical trials have demonstrated improved recurrence-free survival in stage II melanoma patients treated with immune checkpoint inhibitors ^8^.

Interestingly, patients with negative lymph nodes that did not develop distant metastases exhibited lymphangiogenesis and expanded sinusoids associated with plasmablasts. Despite the difference in age between NP and NN patients, this feature was not correlated with patient age. This spatial organization resembles an extrafollicular response (EFR), where plasmablasts are produced from the extrafollicular zone with or without limited germinal center response ^60^. Although we could not directly confirm an EFR in these archival samples, EFR was further supported by a decrease of follicular germinal center cells in the follicle and signs of plasmablast activation outside the germinal centers, including the presence of mast cells in the sinuses^61^. EFRs have been previously linked to autoimmunity^60,70,71^, suggesting that NN patients are perhaps developing plasmablasts with autoreactive antibodies. As such, an intriguing hypothesis emerging from our results is that a successful anti-tumor reaction in the lymph node may require autoimmune-like processes that overcome tolerance barriers. Another possible source of plasmablasts in the extrafollicular region is the reactivation of circulating memory B cells ^72^. Future studies, including spatially resolved immunoglobulin sequencing, could help clarify the origins of these plasmablasts and their role in anti-tumor immunity.

Using spatial data, we predicted development of distant metastases with an area under the curve (AUC) of 79% and 93% for non-metastatic and metastatic LNs, respectively. The classifier was trained solely on protein multiplexed imaging data, and only a subset of the 39 proteins contributed to its predictive power. This suggests that simple, clinically relevant features could improve patient prognostication compared to previously proposed features ^73^. An immediate application for such a model could be in adjusting treatment protocols for melanoma patients with negative sLNs. It was recently shown that immunotherapy can benefit stage IIB and C melanoma patients, having thick or ulcerated primary tumors ^8^. However, since only ±25% of patients with negative LNs will proceed to develop distant metastases within three years of diagnosis^6^, adjuvant treatment of stage II patients may lead to overtreatment with related morbidity and mortality^74^. Our results suggest a potential path to prioritize early-stage high risk patients to treatment.

Importantly, our results were derived from a retrospective cohort that was limited in size and selected to enrich for less common NP and PN patients. Further validation in larger, independent and prospective cohorts is needed. Nevertheless, this study provides a foundation for identifying spatial immune signatures in LNs, which could guide future research on the role of lymph nodes in metastasis prevention and its therapeutic implications.

## Acknowledgements

L.K. holds the Fred and Andrea Fallek President’s Development Chair. She is supported by the Enoch foundation research fund, the Abisch-Frenkel foundation, the Rising Tide foundation, the Sharon Levine Foundation and grants funded by the European Research Council (948811), Israeli Science Foundation (2481/20) and the Rosetrees foundation (10004). M.L. and E.P. are supported by the Dr. Miriam and Sheldon G. Adelson Medical Research Foundation (AMRF). M.L., J.E.C. and L.K. acknowledge the Israel Precision Medicine Partnership program (3830/21). M.L. and L.K acknowledge the Melanoma Research Alliance Team Science Award (1200724). E.P acknowledges the Israel Science Foundation (ISF) Precision Medicine program (1696/20). Y.A. is supported by the CHE/PBC fellowship. I.M. was supported by a EU - Horizon 2020 - MSCA Individual Fellowship (890733).

## Declaration of generative AI and AI-assisted technologies in the writing process

During the preparation of this work the authors used ChatGPT in order to edit and refine the text. After using this service, the authors reviewed and edited the content as needed and take full responsibility for the content of the published article.

## Methods

### Cohort

A retrospective cohort of FFPE sentinel LN biopsies were obtained from 69 melanoma patients. Of the 69 patients, 68 underwent spatial proteomics (MIBI pipeline), and 33 underwent spatial transcriptomics (CosMx pipeline). A full description of the clinical data is available in supplementary Table 1. All patients had a follow-up of at least 5 years, except for one living patient who had a two-year follow-up. In our cohort, no association was observed between treatment and patient groups. (**Fig. S3D).**

### TMA construction

Formalin-fixed, paraffin-embedded (FFPE) tissue blocks of sLNs were retrieved from the tissue archive at the pathology department in Hadassah Medical Center. Regions of interest (2-3) were annotated on corresponding hematoxylin and eosin (H&E)- stained section by a certified pathologist (T.K.H). Next-generation tissue microarrays (TMAs) with 2 mm diameter cores were assembled from these regions using a TMA Grand Master automated tissue microarrayer (3DHistech).

### Antibodies

A summary of antibodies, reporter isotopes, and concentrations can be found in supplementary table 3. Conjugated primary antibodies were purchased from Ionpath or conjugated in-house using the Maxpar X8 Antibody Labeling Kit (Fluidigm, California, USA) or MIBItag Conjugation Kit (Ionpath) according to the manufacturer’s recommended protocols.

### MIBI-TOF staining

Tissue sections (5 µm thick) were cut from the FFPE TMA blocks using a microtome, and mounted on gold-coated slides (Ionpath, Menlo Park, CA) for MIBI-TOF analysis. Slides were baked at 70°C for 20 min. Tissue sections were deparaffinized with 3 washes of fresh xylene. Tissue sections were then rehydrated with successive washes of ethanol 100% (2x), 95% (2x), 80% (1x), 70% (1x), and ultra-pure DNase/RNase-Free water (Bio-Lab, Jerusalem, Israel), (3x). Washes were performed using a Leica ST4020 Linear Stainer (Leica Bio-systems, Wetzlar, Germany) programmed to 3 dips 6 times per wash of 30 seconds. The sections were then immersed in epitope retrieval buffer (Target Retrieval Solution, pH 9, DAKO Agilent, Santa Clara, CA) and incubated at 97°C for 40 min and cooled down to 65°C using Lab vision PT module (Thermo Fisher Scientific, Waltham, MA). After cooling to room temperature, slides were assembled with a coverplate on a sequenza rack and were washed 2 times with 1 ml of TBS-T (Ionpath) diluted in ultrapure water. Sections were then blocked for 1h with 300 µl of 5% (v/v) donkey serum (Sigma-Aldrich, St Louis, MO) diluted in TBS-T. Metal-conjugated antibody mix was prepared in 5% (v/v) donkey serum TBS-T wash buffer and filtered using centrifugal filter, 0.1 µm PVDF membrane (Ultrafree-Mc, Merck Millipore, Tullagreen Carrigtowhill, Ireland). Two panels of antibody mixes were prepared. The first panel contained most of the metal-conjugated antibodies and was incubated overnight at 4°C in a humid chamber. Following overnight incubation, slides were washed 2 times with 1ml TBS-T and once with 300 µl 5% (v/v) donkey serum (Sigma-Aldrich, St Louis, MO) diluted in TBS-T and incubated with the second antibody mix for 1h at room temperature. Slides were washed 3 times with TBS-T and then were removed from the sequenza assembly. Slides were fixed for 5 min in diluted glutaraldehyde solution 2% (Electron Microscopy Sciences, Hatfield, PA) in PBS-low barium. Tissue sections were dehydrated with successive washes of Tris 0.1 M (pH 8.5), (3x), Ultra-pure DNase/RNase-Free water (2x), and ethanol 70% (1x), 80% (1x), 95% (2x), 100% (2x). Slides were immediately dried in a vacuum chamber for at least 1 h prior to imaging.

### MIBI-TOF imaging

Imaging was performed using a MIBIScope mass spectrometer (Ionpath, Menlo Park, CA) with a Xenon ion source. FOVs of 400µm x 400µm or 800µm x 800µm were acquired using a grid of 1024X1024 or 2048X2048 pixels respectively with resolution of 0.45µm, with 1 msec dwell time per pixel. We acquired 193 multiplexed images, 177 from the cohort and 16 control.

### Low level image processing

Multiplexed images were processed to improve image quality including background subtractions, noise removal, and aggregate filtering. The processing was performed using MAUI pipeline as previously described^75^ and further manually denoised with Studio Label^76^.

### CosMx image acquisition and processing

We acquired 77 multiplexed mRNA images using CosMx in two separate runs including two TMAs in each. Image were 3648X5472 or 4480X4480 pixels with a resolution of 0.18µm per pixel, resulting in 848,727 cells. The panel included probes detecting 960 genes, and fluorescent antibodies against: S100B/PMel, CD3, CD45, B2M/CD298 and DAPI. Cells with less than 20 counts were filtered out. A full list of genes can be viewed in supplementary table 2.

### Cell segmentation

The protein data was segmented by DeepCell^22^ (Mesmer’s version 0.11.1 and later was used,) using the dsDNA, MelanA and SOX10 channels as nuclear input and the HLA-I channel as membranal input, resulting in 1,652,274 cells.

The mRNA data was segmented using CellPose^28^ using DAPI as nuclear input and combination of B2M/CD298, S100B/PMel and CD45 as membranal input. Segmentation was performed twice: first, focusing on large tumor cells with S100B/PMel staining, and then on other cells without S100B/PMel staining. Finally, the results of both segmentations were combined. The segmentation of each cell was expanded by 25 pixels during post-processing.

### Cell type classification

The protein data was classified by CellSighter^23^. The training labels were generated by FlowSOM^77^ on 16 images, followed by several rounds of manual cell type correction and annotation. We then trained the model on 12 images and validate on the remaining 4 images. CellSighter was trained to predict 26 cell types. Cell types with less than 50% confidence were labeled as Unidentified. Cells that were predicted as tumor but did not express SOX10 and MelanA were also set to Unidentified.

The mRNA data was classified by InSituType^29^ using the semi-supervised setting. The cell type classification was done on each TMA separately and then merged together by comparing the gene expression of each cell type in each TMA. Results were manually curated following visual inspection of the images.

### Protein expression table

The mean expression of each protein in every cell in the segmentation was calculated by summing the expression of the protein in the cell and dividing by its area in pixels. Values were multiplied by 100, arcsinh transformed, and normalized to the 99th percentile of all positive (greater than 0) cells for that protein. A Cell was considered positive for a specific protein if it had a normalized expression greater than 0.1.

### Gene expression table

The total counts of each gene in every cell in the segmentation was calculated by summing the counts of the gene in the cell. Values were normalized to the 99th percentile of all positive (greater than 0) cells for that gene in each TMA separately. The gene expression in each cell was normalized to a sum of 1.

### T cell phenotyping

We grouped the CD4 T cells and Memory CD4 T cell from the CellSighter’s prediction and clustered them into 20 clusters using K-means clustering based on their functional proteins, including: CD45RA, CD45RO, CCR7, CD69, TIM-3, TCF1-7, PD-1, TOX/2, LAG-3, Ki-67, CD103, GZMB, PD-L1 and IDO-1. To deal with potential spillover of CD45RA, CD4 T cells with positive CD45RA (greater than 1) were set to have zero CD45RA if they had CD20 levels greater than the 1st quantile of CD20 in all B cells for clustering purposes. The optimal number of clusters was determined by examining Silhouette scores and manually validating the quality of the clusters. Each cluster was named based on its average protein expression. Some clusters were manually merged into a single cluster.

A similar analysis was performed on CD8 T cells resulting with 25 clusters. Following manual inspection, two clusters were reclassified as DCs, resulting in 23 clusters.

### Micro environment analysis

A self-supervised deep learning algorithm was developed to identify the microenvironments. The DINO^30^ algorithm was adapted to accept 3D patches, with dimensions H×W×(C+1), as input. H×W represent the dimensions of the patch. C represents the number of cell types. Each channel corresponds to a distinct cell type, where pixel values are 1 for the segmented cells of that type and 0 elsewhere. A binary channel that represents the segmentation of all cells within the patch is included.

DINO is trained to project two different augmentations of the same patch close to each other in the embedding space. To adapt DINO for our data, we added additional augmentations. In total, we randomly applied the following set of augmentations: removing up 3 cells in the patch, switching up the location of up to 5 pairs of cells in the patch, moving the patch, adding random noise, rotating and flipping the patch.

The training included multiple steps starting with training DINO on sliding windows from the images. After training, we projected overlapping sliding window to the embeddings space. We then clustered the embedding space with PhenoGraph^31^ such that, each cluster represents a microenvironment. Next, we trained a logistic regression model to predict the PhenoGraph cluster from the embedding of the patch. For this stage we removed ambiguous clusters from the training. Lastly, for each cell in the data we used DINO to project a patch centered around the cell to the embedding space and used the logistic regression model to predict its cluster, representing a cellular microenvironment.

Microenvironments were grouped to macroenvironments by manually comparing the cell type composition, and spatial proximity. The specific microenvironment definition and grouping can be seen in Figure 2C, D.

Patch sizes for the proteomics data were 64×64 pixels which is equivalent to approximate 29µm and for the transcriptomics data 256×256 pixels which is equivalent to approximate 46µm.

Cells with an assigned microenvironment confidence below 50% were classified as unidentified. Similarly, microenvironments with fewer than 10 cells per field of view (FOV) were also classified as unidentified.

### Macroenvironment segmentation masks

For the proteomics data, macroenvironments that were generated by DINO were processed by removing small areas (5000 pixels), expanding (128 pixels), removing small connected components (area smaller than 15,000 pixels) and expanding again (100 pixels). Each cell was then assigned to a region based on whether it fell within the corresponding region mask. Cells located at the borders between regions may be assigned to multiple regions. Those macroenvironment masks were used in analyses to compare phenotype and cell composition within the macroenvironment.

### Differential gene expression

The Scanpy python module was used to calculate the differentially expressed genes of a specific cell type comparing between the different groups of patients. For each patient, the mean mRNA expression for each gene was calculated across all cell types and macroenvironments. Differentially expressed genes between patient groups were identified by t-test, while controlling for cell type and macroenvironment, genes that were not associated with the cell type of interest were filtered out. In Figure 4K, we excluded dispersed tumors in which the mean distance to the 10 nearest tumor macroenvironment cells exceeded 500 pixels.

### PCA analysis and predictive models

Two models were trained to predict development of distant metastases. For patients with metastatic LNs and for patients with negative LNs. Features that were independently correlated with recurrence, including phenotypes, macroenvironments and protein expression were selected for training. The separation line for the PCA plot was computed using linear Support Vector Machine (SVM) with C=10 and balanced class weights.

A leave-one-out experiment was conducted to assess the predictive power of our selected features. In each iteration, one patient was excluded from the dataset, a model was trained on all remaining patients and then tested on the excluded patient. The model consisted of standard normalization, PCA projection to 3 components and training a SVM classifier with C=1 and balanced class weights. For each patient, the SVM’s prediction and decision function were recorded. These values were used to generate a confusion matrix and ROC curve. The decision function is proportional to the distance of the sample from the SVM’s separating hyperplane.

### Statistic reproducibility

All analyses were done in python (3.11.3). Unless stated otherwise, statistical comparisons were performed using the Mann-Whitney test. For all tests, a minimum number of 5 or 10 cells in the microenvironment, macroenvironment, or cell type lineage was required for a patient to be included in the analysis.

### Figure Illustration

Illustrations were created with BioRender.com.

