## Supplementary Figures for "Immune organization in sentinel lymph nodes of melanoma patients is prognostic of distant metastases"

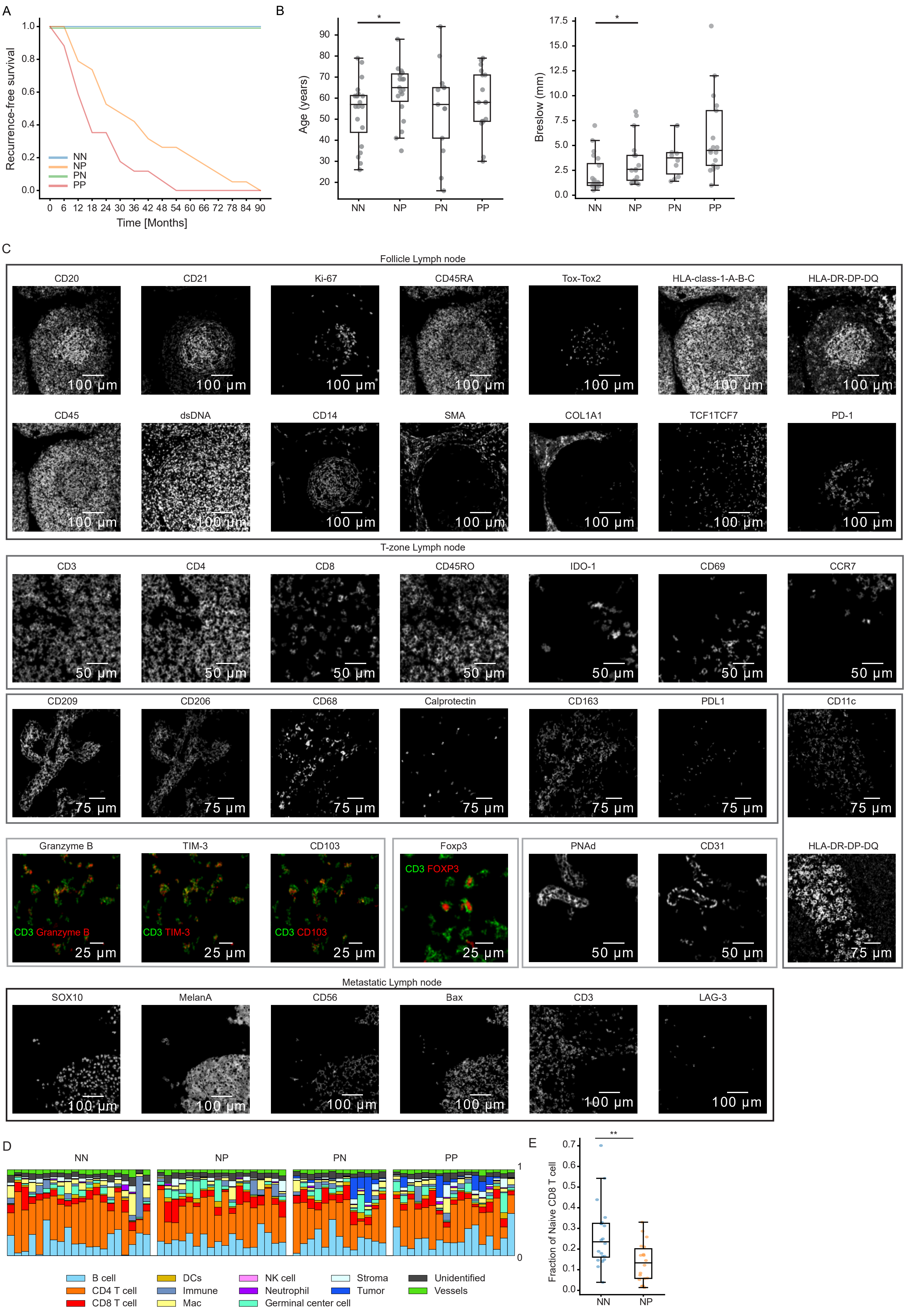

A
Recurrence-free survival
Time [Months]
NN
NP
PN
PP
B
Age (years)
Breslow (mm)
*
C
Follicle Lymph node
CD20
CD21
Ki-67
CD45RA
Tox-Tox2
HLA-class-1-A-B-C
HLA-DR-DP-DQ
CD45
dsDNA
CD14
SMA
COL1A1
TCF1TCF7
PD-1
100 µm
T-zone Lymph node
CD3
CD4
CD8
CD45RO
IDO-1
CD69
CCR7
50 µm
CD209
CD206
CD68
Calprotectin
CD163
PDL1
CD11c
75 µm
Granzyme B
TIM-3
CD103
Foxp3
PNAd
CD31
HLA-DR-DP-DQ
CD3 Granzyme B
CD3 TIM-3
CD3 CD103
CD3 FOXP3
25 µm
Metastatic Lymph node
SOX10
MelanA
CD56
Bax
CD3
LAG-3
D
B cell
CD4 T cell
CD8 T cell
DCs
Immune
Mac
NK cell
Neutrophil
Germinal center cell
Stroma
Tumor
Unidentified
Vessels
E
Fraction of Naive CD8 T cell
**

**Figure S1:**

**(A)** Recurrence-free survival (y-axis) as a function of time (x-axis) for each patient group. NN,NP,PN,PP=20,19,13,17 patients. **(B)** Distribution of ages (y-axis, left) or Breslow scores (right) across patient groups (x-axis). Age analysis: NN,NP,PN,PP=20,19,13,17 patients. Breslow analysis: NN,NP,PN,PP=20,17,10,17 patients. **(C)** Multiplexed images of four FOVs depicting the staining of all antibodies in the panel. For visualization, images were clipped to their 99th percentile and Gaussian blurred. **(D)** Cell type composition (y-axis) for each patient (x-axis) from the proteomics data. NN,NP,PN,PP=20,18,13,17 patients. **(E)** Fraction of Naïve CD8 T cells out of all CD8 T cells (y-axis) in NN and NP patients (x-axis). NN,NP=20,18 patients.

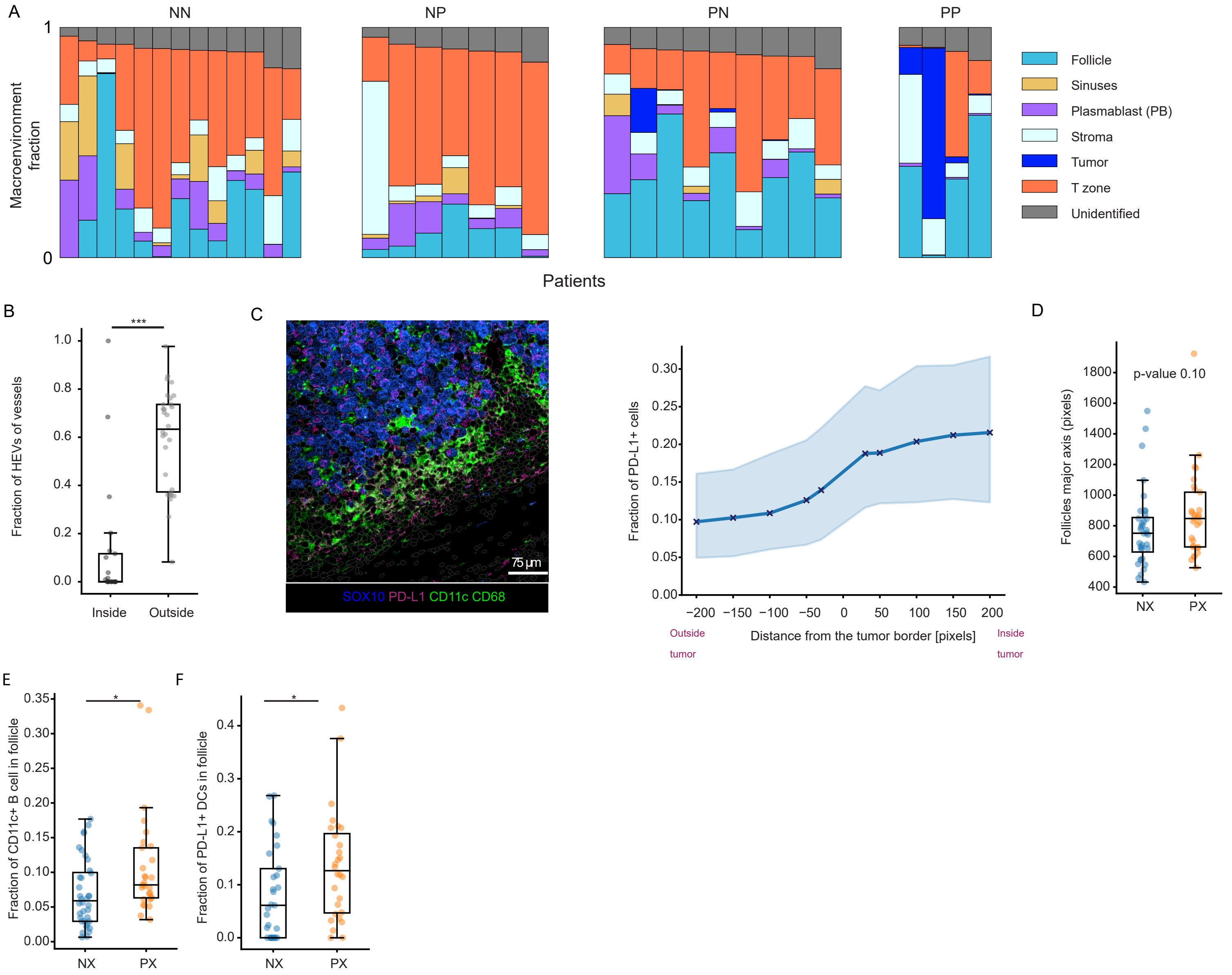


**Figure S2:**

**(A)** Composition of macroenvironments (y-axis) across patients (x-axis) in the transcriptomics data. **(B)** Fraction of HEVs out of all blood vessels (y-axis) inside and outside the metastases (x-axis). n=21 patients inside, 30 outside. **(C)** Left: image of PD-L1 expressed on the tumor. Right: Fraction of PD-L1+ cells (y-axis) as a function of the distance in pixels from the border of the tumor macroenvironment. Maximum of n=21 patients. **(D)** The size of the major axis of follicles (y-axis) in NX and PX patients. NX, PX=36,29 patients. **(E)** Fraction of CD11c+ B cells out of all B cells in the follicle macroenvironment. Only patients with a minimum of 10 B cells were included in the analysis. NX, PX=36,29 patients. **(F)** Fraction of PD-L1+ DCs out of all DCs in the follicle macroenvironment. Only patients with a minimum of 10 DCs were included in the analysis. NX, PX=29,28 patients. **All p-values were calculated using Mann-Whitney statistic test. For (B, E,F) the analysis was limited to regions with more than 100 cells. Follicles regions included germinal centers**.

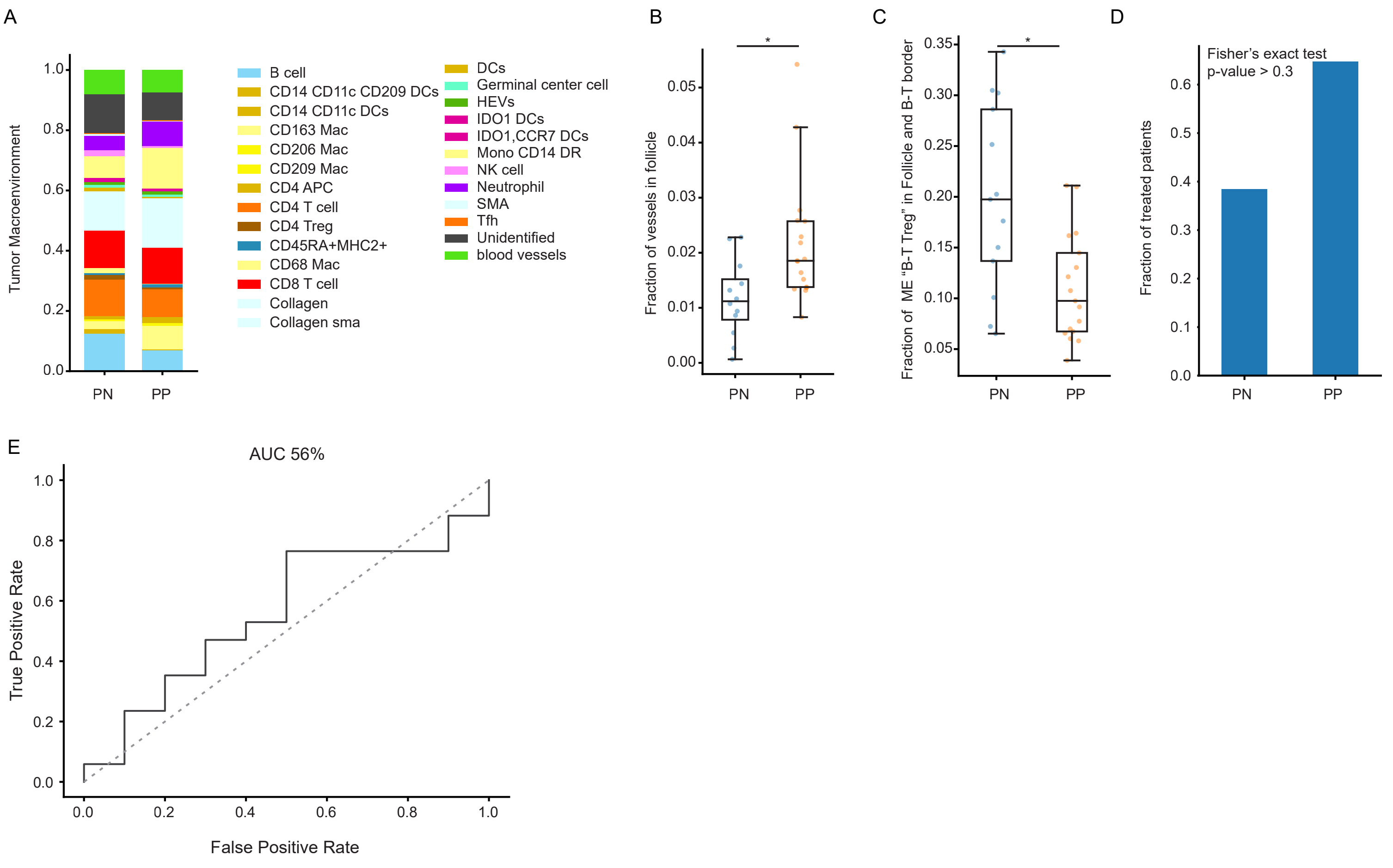


**Figure S3:**

**(A)** Composition of non-tumor cells inside the tumor macroenvironment in PN and PP patients. PN, PP=9, 12 patients. **(B)** Fraction of vessels (y-axis) in the follicles of PN and PP patients (x-axis). Only follicles macroenvironment with more than 100 cells were included in the analysis. PN, PP=12, 17 patients. **(C)** Fraction of the “B-T Treg” microenvironment in the follicles and the B-T zone of PN and PP patients. Only macroenvironment with more than 100 cells were included in the analysis. PN, PP=13, 17 patients. **(D)** Fraction of patients who were treated after diagnosis, and before relapse if relevant, across patient groups.PN, PP=13,17 patients. **(E)** Baseline ROC curve for leave-one-out analysis across patients, using age and Breslow to predict recurrence in positive lymph nodes. PN, PP=10, 17 patients. **All p-values were calculated using Mann-Whitney statistic test. Follicles regions included germinal centers.**

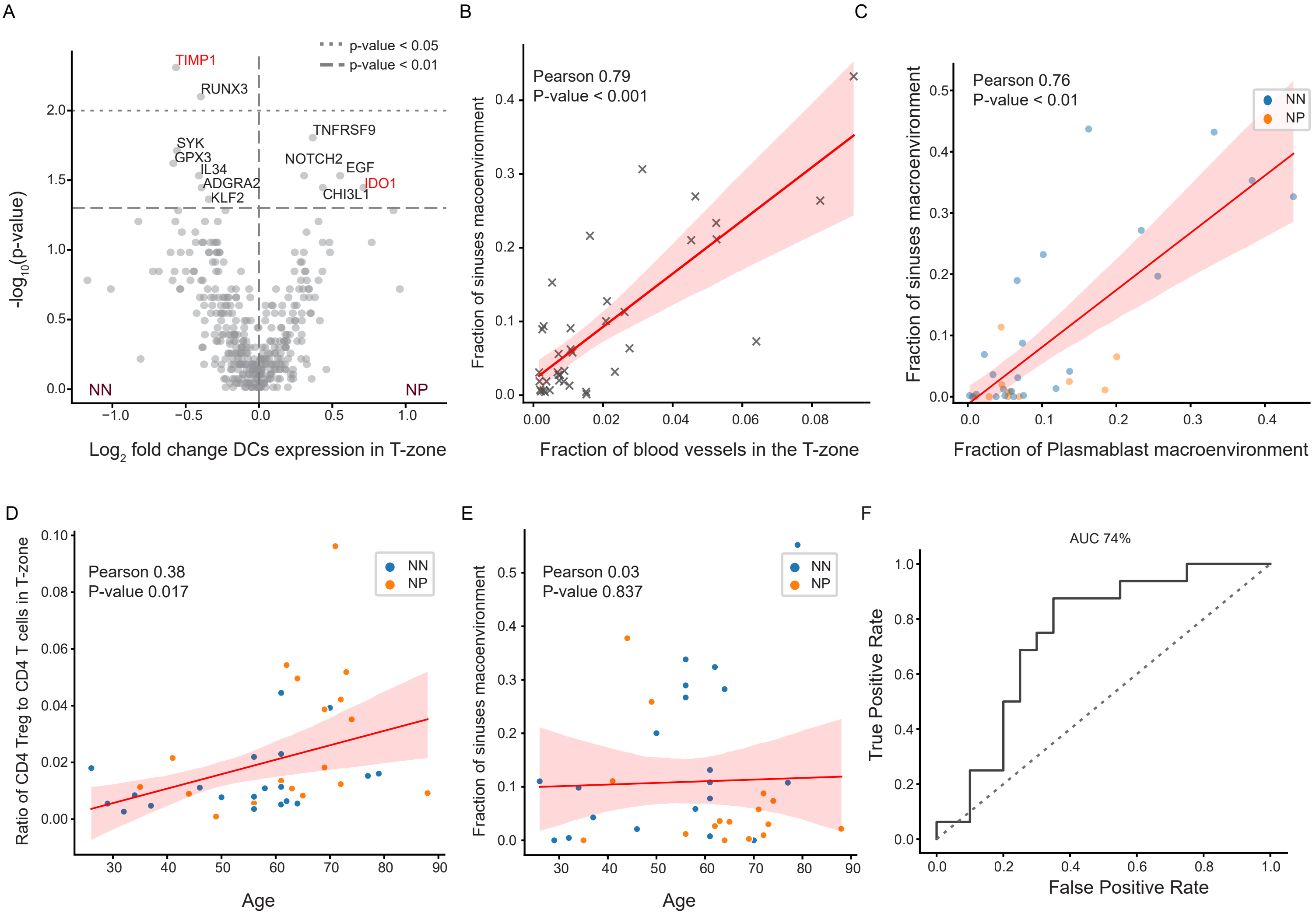


**Figure S4:**

**(A)** Volcano plot showing enrichment of mRNA expression in DCs within the T-zone macroenvironment of NN and NP patients. Dashed lines indicate P-value thresholds of 0.01 and 0.05. NN, NP=13,7 patients. **(B)** Correlation between the fraction of blood vessels in the T-zone (x-axis) and the sinus microenvironment (y-axis). Each dot represents a patient. The analysis was limited to regions with more than 100 cells. n=38 patients. **(C)** Correlation between the plasmablast and sinus macroenvironments in NN (blue) and NP (orange) patients. Each dot represents one field of view. n=34 fovs from 20 patients. **(D)** Correlation between the ratio of Tregs out of all CD4 T cells in the T-zone (y-axis) and patient age (x-axis). NN, P=20,18 patients. **(E)** Correlation between the fraction of the sinus macroenvironment (y-axis) and patient age (x-axis). NN,NP=20,18 patients. **(F)** Baseline ROC curve for leave-one-out analysis across patients, using age and Breslow to predict recurrence in negative lymph nodes. NN,NP=20,16 patients.
